# GIANI: open-source software for automated analysis of 3D microscopy images

**DOI:** 10.1101/2020.10.15.340810

**Authors:** David J. Barry, Claudia Gerri, Donald M. Bell, Rocco D’Antuono, Kathy K. Niakan

## Abstract

The study of cellular and developmental processes in physiologically relevant three-dimensional (3D) systems facilitates an understanding of mechanisms underlying cell fate, disease and injury. While cutting-edge microscopy technologies permit the routine acquisition of 3D datasets, there is currently a limited number of open-source software packages to analyse such images. Here we describe GIANI (djpbarry.github.io/Giani), new software for the analysis of 3D images, implemented as a plugin for the popular FIJI platform. The design primarily facilitates segmentation of nuclei and cells, followed by quantification of morphology and protein expression. GIANI enables routine and reproducible batch-processing of large numbers of images and also comes with scripting and command line tools, allowing users to incorporate its functionality into their own scripts and also run GIANI on a high-performance computing cluster. We demonstrate the utility of GIANI by quantifying cell morphology and protein expression in confocal images of mouse early embryos and by segmenting nuclei from light sheet microscopy images of the flour beetle embryo. We also validate the performance of the software using simulated data. More generally, we anticipate that GIANI will be a useful tool for researchers in a variety of biomedical fields.

## Introduction

The ability to routinely acquire multi-dimensional datasets with modern microscopy techniques is transforming, among other fields, cell biology, developmental biology and cancer research. There has long been an acceptance that two-dimensional (2D) cell cultures may not accurately recreate behaviours found in complex, three-dimensional (3D) *in vivo* environments [1]. Commonly-used 3D culture formats include, but are not limited to, populations of single cells in organotypic matrices, spheroid models, tissue sections or whole embryos and organisms.

However, the development of software for the quantitative analysis of such data has not kept pace with imaging advances and there is now a pressing need for automated solutions [2]. Manual annotation of such data is not feasible, due to the time required to do so. Commercial packages, such as Imaris (Bitplane) and Vison4D (Arivis) provide excellent visualisation functionality and are also equipped with analysis tools. However, licenses for such software are expensive and they also rely on proprietary file formats. Furthermore, the closed-source nature of such software prevents detailed interrogation of specific calculations and processes.

There are numerous excellent, freely-available bioimage analysis tools in the open source domain. However, their support for 3D analysis is either limited (for example, several CellProfiler [3] modules are not yet compatible with 3D images) or challenging to execute and/or automate for the uninitiated (FIJI [4], Icy [5]). This can lead to undesirable compromises being made, such as 2D slices from a 3D volume being analysed individually, blinding the analysis to information in adjacent slices. Alternatively, 3D data may be compressed into 2D via projection, which, consequently, artificially reduces distances between objects and can lead to spurious results. A number of very useful open-source MATLAB (MathWorks, Cambridge, UK) -based tools have also been implemented, most notably MINS, which has been effectively used to analyse embryo datasets [6] and LOBSTER [7]. However, these require the purchase of a MATLAB license.

We have therefore developed GIANI (**G**eneral **I**mage **A**nalysis of **N**uclei-based **I**mages; https://djpbarry.github.io/Giani), a generally-applicable, open source tool, implemented as a plugin for the widely-used image analysis platform, FIJI. With an emphasis on detection and segmentation of cells in 3D microscopy images, GIANI has been implemented specifically with batch-processing in mind. While an understanding of fundamental concepts of bioimage analysis is beneficial, GIANI’s user interface has been implemented in a wizard format to facilitate use by non-specialists and is fully documented (https://github.com/djpbarry/Giani/wiki). Analysis protocols may be reproduced by loading a single, small parameter file.

The utility of GIANI is illustrated here using three examples. In the first, we generated a series of simulated datasets to evaluate the accuracy of segmentations produced by GIANI using known “ground truths”. In the second proof-of-concept we use a series of mouse preimplantation embryo datasets and demonstrate the ability of GIANI to detect variations in morphology and protein expression in different experimental conditions. Finally, to show that GIANI can also be used on much larger datasets, we present segmentations of nuclei from light sheet microscopy images of the flour beetle embryo. It should be noted that GIANI can be used to quantify 3D images from a range of cellular and developmental contexts and we anticipate that it will be a useful tool to automate quantification of a wide variety of complex imaging data.

## Materials and methods

### Simulated data generation

Simulated data sets were generated using an extension of a previously described method [8]. Simulated nuclei positions were generated as previously described [8], then cell membranes were approximated using a Euclidean Distance Map constructed around nuclei. The simulated images were then convolved with a Gaussian point-spread function, sub-sampled and noise added from a gamma distribution. The complete code for simulated image generation is available on GitHub (https://github.com/djpbarry/Embryo-Generator).

### Metrics for segmentation quality assessment

We calculated the cell count error *E*_*c*_, as

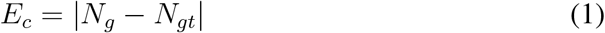

where *N*_*gt*_ is the actual number of cells present in a simulated embryo and *N*_*g*_ is the number counted by GIANI. Nuclear (*E*_*nl*_) and cell (*E*_*cl*_) centroid localisation errors were calculated based on the euclidean distance between the known ground truth centroids and the centroids of the segmentations produced by GIANI.

### Mouse zygote collection

Four- to eight-week-old (C57BL6 × CBA) F1 female mice were super-ovulated using injection of 5 IU of pregnant mare serum gonadotrophin (PMSG; Sigma-Aldrich). Forty-eight hours after PMSG injection, 5 IU of human chorionic gonadotrophin (HCG; Sigma-Aldrich) was administered. Superovulated females were set up for mating with eight-week-old or older (C57BL6 × CBA) F1 males. Mice were maintained on a 12 h light–dark cycle. Mouse zygotes were isolated in EmbryoMax FHM mouse embryo media (Sigma-Aldrich; MR-122-D) under mineral oil (Origio; ART-4008-5P) and cumulus cells were removed with hyaluronidase (Sigma-Aldrich; H4272). All animal research was performed in compliance with the UK Home Office Licence Number 70/8560.

### Mouse embryo culture

Mouse embryos were cultured in drops of pre-equilibrated Global medium (LifeGlobal; LGGG-20) supplemented with 5 mg/ml protein supplement (LifeGlobal; LGPS-605) and overlaid with mineral oil (Origio; ART-4008-5P). Preimplantation embryos were incubated at 37°C and 5.5% CO_2_ and cultured up to the day of analysis.

### Inhibitor treatment

Inhibitor experiment was performed as previously described [9]. Briefly, the aPKC inhibitor CRT0276121 (Cancer Research Technology LTD) was dissolved in DMSO to 10 mM stock concentration and diluted to the optimal concentration of 8 *µ*M in pre-equilibrated embryo culture media. Mouse embryos were incubated in pre-equilibrated media with 8 *µ*M of CRT0276121 from 4-cell to morula stage. Control mouse embryos were developed in pre-equilibrated media where volume-matched DMSO was added.

### Immunofluorescence

Embryos were fixed with freshly prepared 4% paraformaldehyde in PBS that was pre-chilled at 4°C. Embryo fixation was performed for 20 min at RT and then the embryos were transferred through 3 washes of 1X PBS with 0.1% Tween-20 to remove residual paraformaldehyde. Embryos were permeabilized with 1X PBS with 0.5% Triton X-100 and then blocked in blocking solution (3% BSA in 1X PBS with 0.2% Triton X-100) for 2 h at RT on a rotating shaker. Then, embryos were incubated with primary antibodies (listed in Table 1) diluted in blocking solution overnight at 4°C on rotating shaker.

**Table 1.** Primary antibodies used in this study and the dilution at which they were used.

| Antibody | Supplier | Catalogue Number | Dilution |
| --- | --- | --- | --- |
| YAP1 | Abnova | H00010413-M01 | 1:50 |
| GATA3 | R&D | AF2605 | 1:200 |
| E-CADHERIN | Life Technologies | 131900 | 1:400 |

The following day, embryos were washed in 1X PBS with 0.2% Triton X-100 for 20 min at RT on rotating shaker and then incubated with secondary antibodies diluted in blocking solution for 1 h at RT on rotating shaker in the dark. Next, embryos were washed in 1X PBS with 0.2% Triton X-100 for 20 min at RT on rotating shaker. Finally, embryos were placed in 1X PBS with 0.1% Tween-20 with Vectashield with DAPI mounting medium (Vector Lab; H-1200) (1:30 dilution). Embryos were placed on *µ*-Slide 8 well dishes (Ibidi; 80826) for confocal imaging.

### Image Acquisition

All images were acquired on a Leica SP5 laser scanning confocal microscope using a Leica 1.3 NA 63x HCX PL APO CS glycerol objective and a voxel size of approximately 0.1 × 0.1 × 1.0 *µ*m in x, y and z, respectively.

### Software

GIANI was written using Java 8 as a plug-in for FIJI [4], making extensive use of the underlying FIJI and ImageJ [10] libraries. A number of other open-source projects are leveraged. Reading of image data is facilitated by interfacing to Bio-Formats [11], making GIANI compatible with a wide range of file formats (https://docs.openmicroscopy.org/bio-formats/6.7.0/supported-formats.html). Detection of nuclear blobs makes use of either TrackMate’s spot detector [12] or FeatureJ (https://imagescience.org/meijering/software/featurej). Segmentation of cells and nuclei takes advantage of the marker-controlled watershed functionality in MorphoLibJ [13] and 3D Image Suite [14]. The browsing of results is based upon the 3D Region of Interest (ROI) Manager from 3D Image Suite. Complete source code, documentation and test data are available online (https://djpbarry.github.io/Giani).

### Statistical Analysis

All statistical analyses in this study were performed using RStudio (www.rstudio.com). The tests performed in this paper are two-sample Wilcoxon tests (also know as Mann-Whitney). All box plots show the median and inter-quartile range, with the whiskers extending 1.5 times the inter-quartile range from the 25th and 75th percentiles. Unless otherwise stated, each dot in dot plots represents a single cell.

## Results & Discussion

### Software implementation

The design philosophy behind GIANI is inspired by CellProfiler. It is assumed that the user wishes to detect “primary objects” of some sort (typically cell nuclei), followed by the subsequent segmentation of “secondary objects” (typically cells) and then wishes to measure either the morphology of said objects, or the expression of a fluorescent signal within these objects. The principal difference in the case of GIANI is that, in order to facilitate ease of use, the order of steps in the pipeline is fixed, although some flexibility is present where necessary (Fig 1).

**Fig 1.**
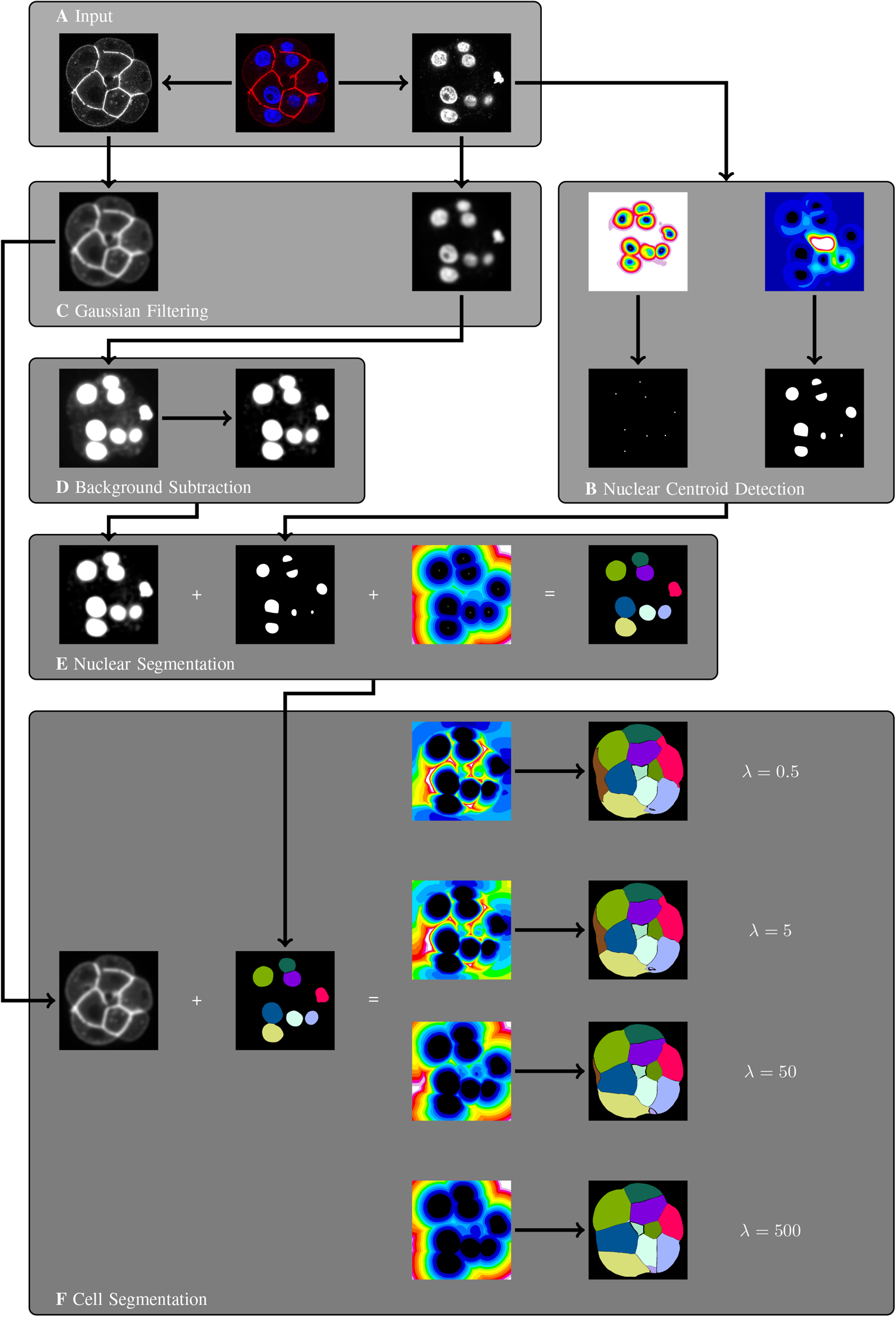
Overview of the algorithms underpinning GIANI. All images represent a single slice of a 3D stack and are for illustrative purposes only. A: An example dataset, consisting of a mouse embryo showing DAPI (blue) and E-Cadherin (red). GIANI can accept as input any image data that is readable by Bio-Formats [11]. B: Nuclear are first approximated using one of two blob detectors - Laplacian of Gaussian (left) or Hessian (right). C: Gaussian filtering is used to suppress noise in the channels used for nuclear and cell segmentation. D: The nuclear channel is then subjected to top-hat filtering to remove background - contrast has been increased to illustrate the effect of the filter. E: Nuclear segmentation is achieved using a marker-controlled watershed approach, with the background-subtracted image from D serving as the input and centroids from B serving as the seeds. F: Cell segmentation is achieved using the same marker-controlled watershed approach, with the filtered image from C serving as the input and nuclei segmentations from E serving as the seeds.

#### File Formats

GIANI can accept as input any images that are readable by Bio-Formats [11], which, at the time of writing, can read almost 160 different formats. It is expected that this data will be three-dimensional in nature, containing any number of different channels. As a minimum requirement, in order to facilitate accurate segmentation, these channels must contain some form of marker for both nuclei and cells. The nuclear marker must be a volume marker, while the cell marker can be either membrane-localised or volumetric in nature, although the former is preferred for more accurate segmentation. In the example pipeline illustrated in Fig 1, a nuclear volume marker (DAPI) and cell membrane marker (E-Cadherin) are used.

#### Detection of Nuclear Centres

The nuclear channel is first subjected to one of two forms of blob detection to estimate the centroid location of each nucleus (Fig 1B). The first option is Laplacian of Gaussian (LoG) blob detection, which involves the application of an LoG filter to the image and then identifying local extrema (Fig 1B, left panels). This is implemented in GIANI by using the spot detector of TrackMate [12], which is based on LoG detection. The advantage of this approach is that it is relatively fast and generic. The disadvantage is it assumes the objects to be detected are approximately Gaussian in nature, so will likely perform poorly for irregularly shaped nuclei.

For this reason, a second option is given for nuclear centroid detection, based on the eigenvalues of the image’s Hessian matrix (Fig 1B, right panels) - this is similar to the approach used for nuclear detection in MINS [6]. The Hessian is a square matrix of second-order partial derivatives describing the local curvature in the image. For a detailed explanation on how Hessian eigenvalues can be used to detect blobs in images, see [15]. This is implemented in GIANI by using FeatureJ, a sub-component of the ImageScience plugin (https://imagescience.org/meijering/software/featurej). This approach has the advantage of being capable of detecting “blobs” of any shape, such as irregularly shaped nuclei. The main disadvantage is that it is much more computationally demanding than the simpler LoG detector, so will take longer to run.

#### Suppression of Noise and Background

Prior to full segmentation of nuclei, Gaussian filtering is employed to smooth any noise that may have been present in the input image (Fig 1C). To both enhance the nuclei and homogenise the image background, the output of the Gaussian filtering operation is input into a top-hat filter (Fig 1D), implemented using MorphoLibJ [13]. Top-hat filtering can be computationally expensive for large datasets, so GIANI includes an option to downsize datasets by a specified factor prior to this step - the dataset is then restored to its original size post-filtering.

#### Segmentation of Nuclei

Nuclei are fully segmented using a marker-controlled watershed approach (Fig 1E). GIANI uses MorphoLibJ to achieve this, with the previously-detected nuclear centroids serving as seeds and the Euclidean Distance Transform of the centroid mask, calculated using the 3D ImageJ Suite [14], serving as the image to be flooded. In addition, the top-hat filtered nuclei channel is thresholded (using one of FIJI’s in-built thresholding algorithms, specified by the user) to act as a mask, restricting the overall extent of the segmentation.

#### Segmentation of Cells

Cells are segmented using the same marker-controlled watershed approach (Fig 1F), with the previously-segmented nuclei serving as seeds and the distance transform of the nuclei serving as the image to be flooded. The Gaussian-filtered cell channel (Fig 1C) is thresholded to act as a mask in this case. The nature of the distance transform used is dependent on whether a membrane marker or volume marker is used to identify the cells. In the case of a volume marker, a standard Euclidean Distance Transform is used. In the case of a membrane marker, a modified distance transform is used, similar to that described by [16], where the distance between adjacent voxels, *d*, is calculated based on a Riemannian metric defined in terms of the image *I* and a regularisation parameter, *λ*:

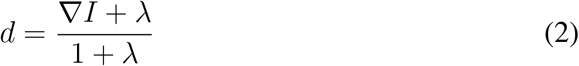

It can be seen that, as *lim*_*λ*→∞_, *d* tends towards the Euclidean distance.

### Analysis of simulated embryos

To first validate the performance of GIANI, we generated a series of 189 “simulated embryos” exhibiting different levels of signal-to-noise ratio and cell density (Fig 2A). Simulated data has the significant advantage of having a known “ground truth” (Fig 2B). That is, because the images are generated artificially, we know what the “correct” segmentation should look like. This permits us to compare the segmentation results generated with any piece of analysis software with the known “true” values.

**Fig 2.**
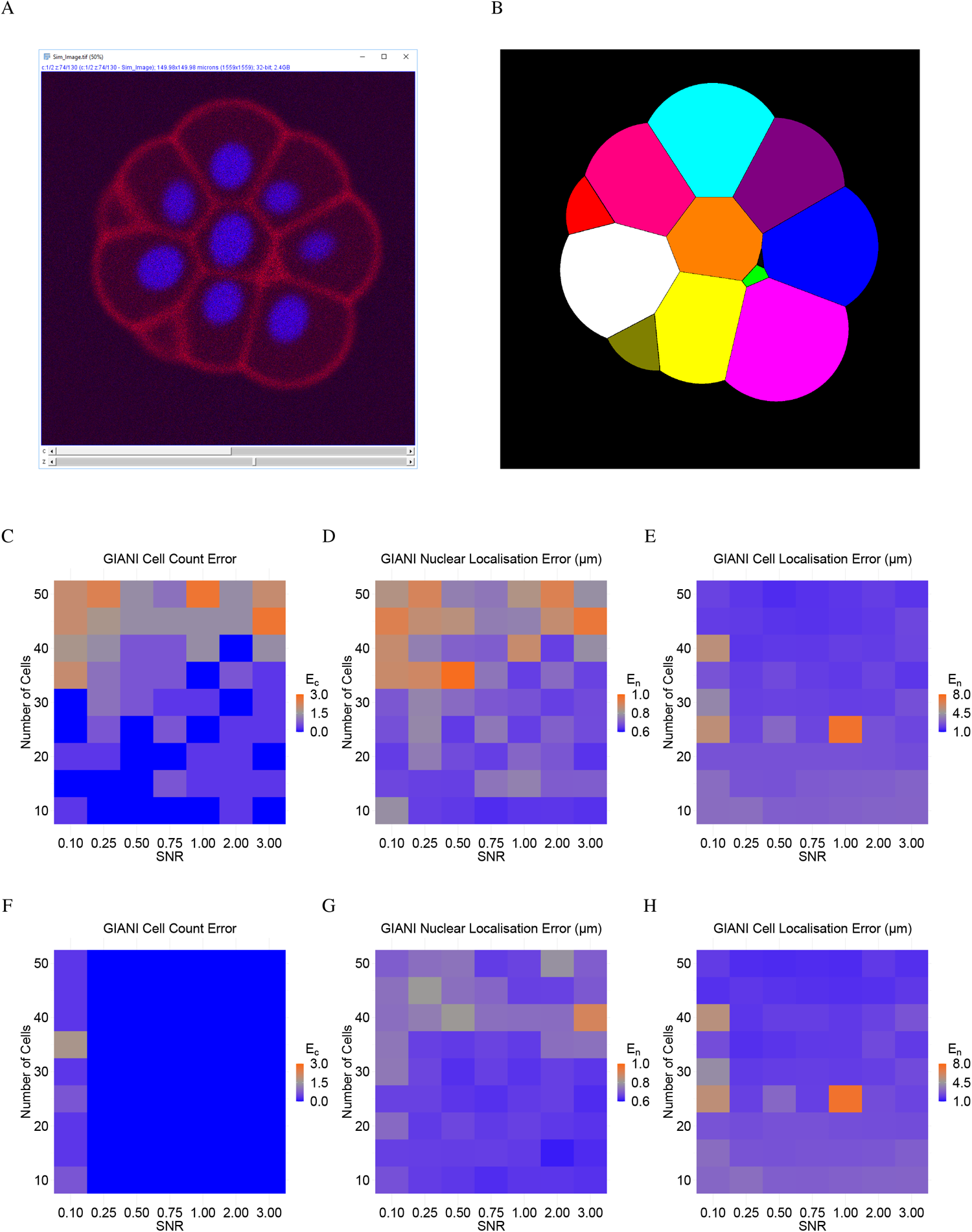
Validation of GIANI using simulated embryo data. In each of the heat maps, a single tile represents the average of three simulated embryos. The GIANI settings used to produce this data are provided as File S3. A: A 2D slice of an exemplar 3D simulated embryo. B: The ground truth segmentation of ‘A’. C - H: Absolute errors in cell counts (*E*_*c*_, calculated according to Eq 1), nuclear centroid localisation error (*E*_*nl*_) and cell centroid localisation error (*E*_*cl*_) produced by GIANI for simulated embryos with the indicated number of cells and signal-to-noise ratios (SNR). Results were obtained using either GIANI’s basic (C - E) or advanced (F - H) nuclear detector.

The number of false positives and false negatives produced by GIANI is consistently extremely low, although cell count errors do generally increase with cell density and decreasing signal-to-noise ratio (Fig 2C, F). At high signal-to-noise ratios and low cell densities, GIANI localises nuclei to within approximately 600 nm of their true positions (Fig 2D, G). Even at high cell densities and low signal-to-noise ratios, this error never exceeded 1 *µ*m (the simulated nuclei are ellipsoidal with axis dimensions of approximately 10.0 × 7.5 × 7.5 *µ*m). In addition to nuclear localisation error, we also quantified the cell centroid localisation error on a per cell basis (Fig 2E, H).

To benchmark GIANI against other available software, we analysed the same simulated embryos using Imaris (Fig S1). This demonstrated that the performance of GIANI is comparable to Imaris when it comes to nuclear localisation (Fig S1 B, D) and that GIANI is superior in detecting the correct number of cells (Fig S1 A, C). We also performed a limited evaluation of CellProfiler on a subset of the same simulated data, which resulted in errors far larger than those produced by GIANI (Fig S2).

The accurate quantitative analysis of a multi-cellular image dataset, such as an embryo, is dependent on the correct identification and segmentation of nuclei. While GIANI presently employs generic blob detectors for this purpose, which is similar to other previously-described methods [6], other approaches could be incorporated [17–21]. In particular, the ability to load pre-trained neural networks, by leveraging DeepImageJ [22], for example, or machine-learning classifiers trained with, for example, Weka [23] or Ilastik [24], may be added in a future release.

In addition to the convolution of the ground truth data, the simulation process also takes into consideration depth of slices within a sample when modelling fluorescent intensity - deeper slices will be more prone to scattering effects and therefor exhibit lower intensity. This means that finding a single global thresholding strategy that does not under-segment the dimmer cells while also not over-segmenting the brighter cells is challenging. Taking all this into consideration, any metric used to assess the accuracy of cell segmentations will have its limitations.

There are strategies that could be employed to mitigate against these factors. For example, some form of adaptive thresholding, whereby the intensity threshold changes according to z-location, could be used. However, while an optimal adaptive thresholding strategy could be found for a given dataset (such as the simulated data used in this study), implementing a universally-applicable strategy would be difficult. One of the principle design aims of GIANI was simplicity of use, which does not allow for the incorporation of a variety of case-specific segmentation approaches.

### Analysis of mouse embryos

We subsequently applied GIANI to the analysis of two populations of mouse preimplantation embryos at the morula stage, one control (*n* = 18) and one treated with a small molecule inhibitor [9] (*n* = 20). At this stage, two distinct cell populations are discernible: inner and outer cells. In subsequent cell divisions, a blastocyst is formed, whereby the inner cells give rise to an inner cell mass (ICM), and the outer cells become the trophectoderm (TE), a polarized epithelium that will form fetal components of the placenta [25].

At the morula stage, inner and outer cells display different polarisation states, which influence their cell fate acquisition. The outer cells acquire an apical domain, enriched with the atypical protein kinase C (aPKC) [26]. In the polar outer cells, aPKC prevents activation of downstream Hippo pathway kinases, Large tumor suppressor kinases 1/2 (LATS1/2) [27]. Consequently, in outer cells, YAP1 accumulates in the nucleus, where it promotes the expression of GATA3 [28]. In contrast, in the apolar inner cells, the activation of the Hippo pathway results in YAP1 cytoplasmic retention, thus maintaining the inner cells in an unspecified state [27, 29, 30].

We therefore divided cells within each embryo into an ‘inner’ and ‘outer’ population (Fig 3A, B). This division is based on the distance of the detected nuclear centroid from the embryo centroid. A cell was classified as ‘inner’ if the following condition held true:

**Fig 3.**
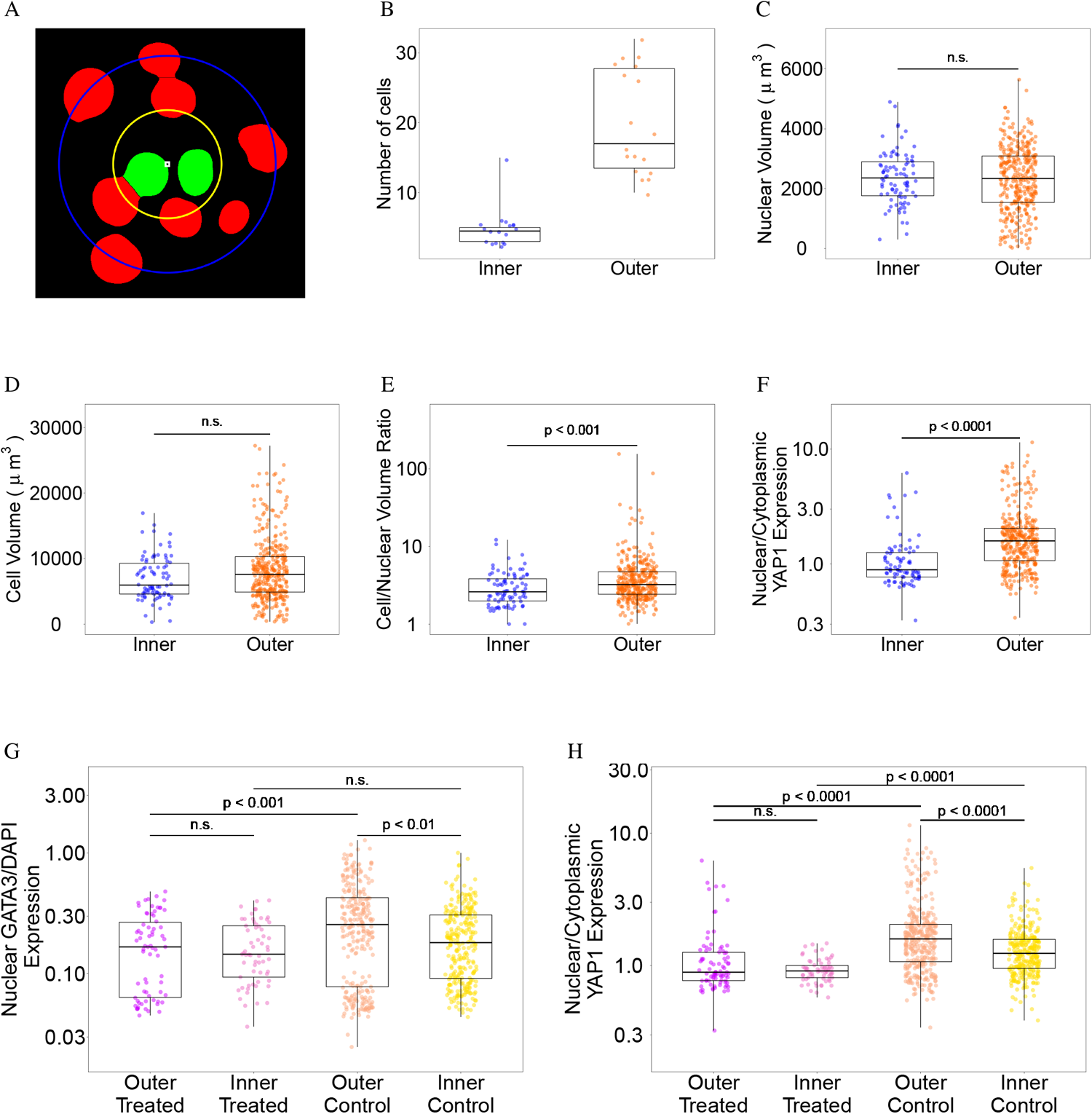
GIANI reveals differences in morphology and protein expression in mouse embryos. Unless otherwise stated, each dot represents a single cell. The GIANI settings used to produce this data are provided as File S4. A. Illustration of the division of embryo cells into ‘outer’ (red) and ‘inner’ (green) sub-populations. The embryo centroid is indicated by the white square. The blue circle has radius *D*_*m*_ (see Eq 3) and indicates the distance from the embryo centroid to the most distant nucleus centroid. The radius of the yellow circle is *D*_*T*_ × *D*_*m*_ (Eq 3). B. The number of cells in each embryo divided into outer and inner sub-populations using a value of 0.5 for *D*_*T*_ in control embryos. Each dot represents a single embryo (*n*_*control*_ = 18). C: Volume of nuclei in inner/outer populations in control embryos. D: Volume of cells in inner/outer populations in control embryos. E: Ratio of cell-to-nuclear volume in inner and outer cells in control embryos. F: Nuclear/cytoplasmic ratio of YAP1 expression in control embryos. G: Difference in expression profiles of nuclear GATA3 expression, normalised to DAPI, in control and treated embryos (*n*_*treated*_ = 20). H: Difference in nuclear/cytoplasmic ratiometric expression profiles of YAP1 in control and treated embryos.

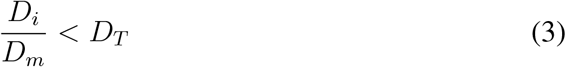

where *D*_*i*_ is the Euclidean distance of cell *i* from the embryo centroid, *D*_*m*_ is the maximum Euclidean distance from any cell in the same embryo to the embryo centroid and *D*_*T*_ is an arbitrarily set distance threshold. For the purposes of this study, we chose a value of 0.5 for *D*_*T*_ - it was found that modifying this value slightly (±10%) did not significantly impact the results (data not shown). The embryo centroid was calculated as the average of all nuclear centroids.

### GIANI shows differences in cell morphology, YAP1 and GATA3 expression within inner and outer cells in control embryos

We then applied GIANI to detect variations in morphology and protein expression in mouse preimplantation embryos. We began with a morphological analysis of cells in control embryos. No significant difference was found between median nuclear volumes in inner and outer cell populations (Fig 3C; *p* = 0.861, Wilcoxon rank sum test). A difference in median cell volume between the two populations was also identified, although it fell slightly short of being statistically significant (Fig 3D; *p* = 0.069). However, a comparison of cell-to-nucleus volume ratios revealed a significant difference between median values, with outer cells having a proportionately greater cytoplasmic volume than inner cells (Fig 3E; *p* < 0.001). We also confirmed previous analysis of YAP1 translocation [31], illustrating that nuclear localisation is significantly higher in outer versus inner cells (Fig 3F; *p* < 0.001). Similarly, GATA3 expression was shown to be higher in outer compared to inner cells (Fig 3G; *p* < 0.0001).

### GIANI reveals differences in YAP1 and GATA3 expression in mouse embryos after pharmacological treatment

To further demonstrate the utility of GIANI, we sought to analyse GATA3 and YAP1 expression after treating mouse embryos with a small molecular inhibitor against aPKC, the upstream regulator of YAP1 and GATA3. The aPKC inhibitor, CRT0276121, has previously been confirmed to specifically inhibit aPKC in various biological and cellular contexts [32–34]. Specifically, in mouse pre-implantation embryos, aPKC inhibition has been recently shown to efficiently abrogate YAP1 and GATA3 expression in outer cells [9].

Expression of GATA3 in the nucleus (normalised to DAPI to correct for diminished signal intensity with increasing sample depth) was found to be similarly low in the inner cells of both control and treated embryos (Fig 3G; *p* = 0.841). However, GATA3 expression in outer cells was found to be significantly lower in the treated embryos relative to the control group (*p* < 0.002). Normalised expression of GATA3 was also found to be significantly higher in outer cells of control embryos relative to their inner cells (*p* < 0.001), while the same was not true of treated embryos (*p* = 0.171). In addition, differences in distribution are evident between control and treated embryos, with two distinct populations evident in both inner and outer cells in control populations (Fig 3G).

While no statistically significant difference in nuclear YAP1 expression (normalised to cytoplasmic expression) between control and treated cells were observed when comparing inner cells (Fig 3H; *p* = 0.609), a large difference was observed in outer cells (*p* < 0.0001). However, nuclear/cytoplasmic YAP1 expression was still higher in the outer cells of treated embryos versus inner (*p* < 0.0001). Altogether, this demonstrates that GIANI allows for the automated quantification of expression differences between cells following perturbation.

### Analysis of blastocysts

Finally, to illustrate that GIANI can be run successfully on later stage mouse embryos, we analysed two examples of blastocysts and compared with Imaris. While the resultant segmentations are less than perfect, a qualitative assessment (in the absence of ground truth segmentations) shows that the results produced by GIANI are at least comparable to those obtained using Imaris (Fig S3).

### Analysis of larger datasets

#### Analysis of large simulated dataset

To test GIANI’s ability to handle more complex datasets, we generated a large simulated volume consisting of approximately 2,000 nuclei (File S1). We compared the segmentations generated by GIANI with those generated by Imaris (Fig 4). A qualitative assessment shows that while both software successfully segment the vast majority of cells, errors are apparent, particularly in cases were nuclei are highly clustered (Fig 4A). We attempted to quantify these success rates using a number of different metrics.

**Fig 4.**
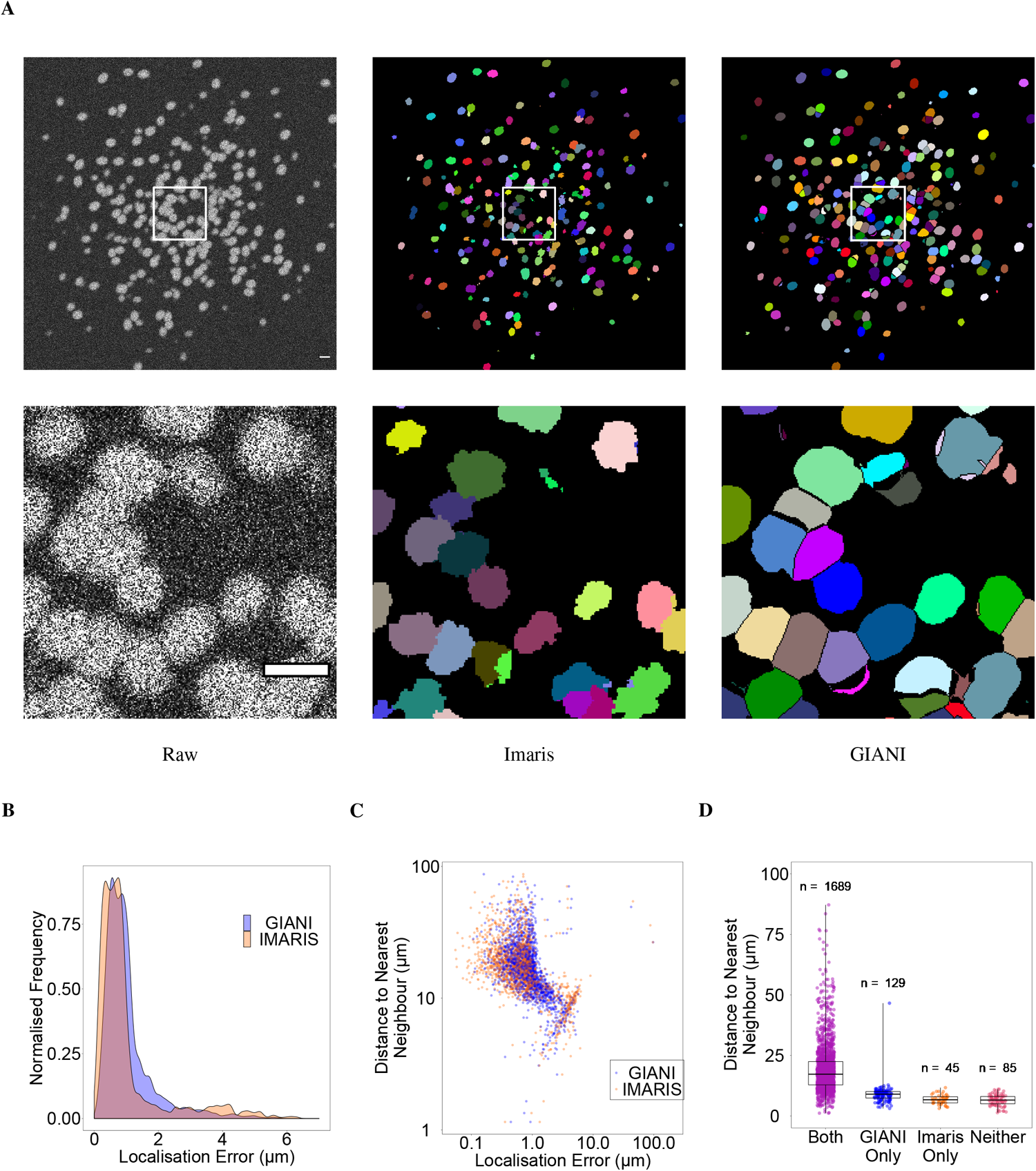
Demonstration of GIANI with a large simulated dataset. The GIANI and Imaris settings used to produce this data are provided as File S5. A: Illustration of the segmentations produced by GIANI and Imaris on a large simulated dataset (File S1). The top row shows a single slice of each 3D volume, while the bottom row shows the magnified views of the boxes in the top row images. Scale bars are all equivalent to 20 *µ*m. B: Distribution of localisation errors produced by both Imaris and GIANI in detecting the simulated nuclei in File S1. C: Relationship between localisation error for detected nuclei and the distance of each nucleus to its nearest neighbour. D: Influence of distance of nuclei to their nearest neighbour on successful detection.

We first compared the distribution of localisation errors produced by GIANI and Imaris (Fig 4B). While in both cases nuclei were localised to within approximately 1 *µ*m or less of their true locations most of the time, the localisation errors produced by Imaris were, overall, marginally lower. But, Imaris had a greater tendency to produce outliers (localisation errors above approximately 2*µ*m).

We next looked for any link between localisation error and nuclear density. To do so, we examined the relationship between the localisation error for each successfully detected nucleus and the distance between that nucleus and its nearest neighbour (calculated as the distance between nuclear centroids; Fig 4C). While the relationship between these two variables is highly non-linear, it is apparent that, for both GIANI and Imaris, a trend exists - localisation error increases as the distance between nuclei decreases.

Finally, we compared the success rates of GIANI and Imaris in detecting the simulated nuclei and investigated whether nuclear density influenced the likelihood of successful nuclear detection (Fig 4D). We found that while both GIANI and Imaris successfully detected the vast majority of nuclei, the success rate was slightly higher for GIANI (93.3% versus 89.0%). There was also a clear relationship between inter-nuclear distance and likelihood of successful detection. Of the 85 nuclei that both Imaris and GIANI failed to detect, 95% were 10 *µ*m or less from their nearest neighbour. Given that the longest dimension of the simulated nuclei is approximately 10 *µ*m, this is perhaps not an unreasonable result - two such nuclei that are less than 10 *µ*m apart at their centres are likely to significantly overlap.

#### Analysis of a dataset derived from light sheet microscopy

To further emphasise the capacity of GIANI to analyse large volumes of data, we tested it on a *Tribolium castaneum* embryo dataset derived from light sheet microscopy (File S2), again benchmarking against Imaris (Fig 5). We obtained this data from the Cell Tracking Challenge (http://celltrackingchallenge.net/3d-datasets) and, unfortunately, the ground truth data available is very limited (annotations for only three cells is provided). However, the intention here is to demonstrate that GIANI is capable of segmenting large numbers of nuclei in such data (approximately 5,600 were detected) and the results are comparable to those obtained using market-leading commercial software (Imaris).

**Fig 5.**
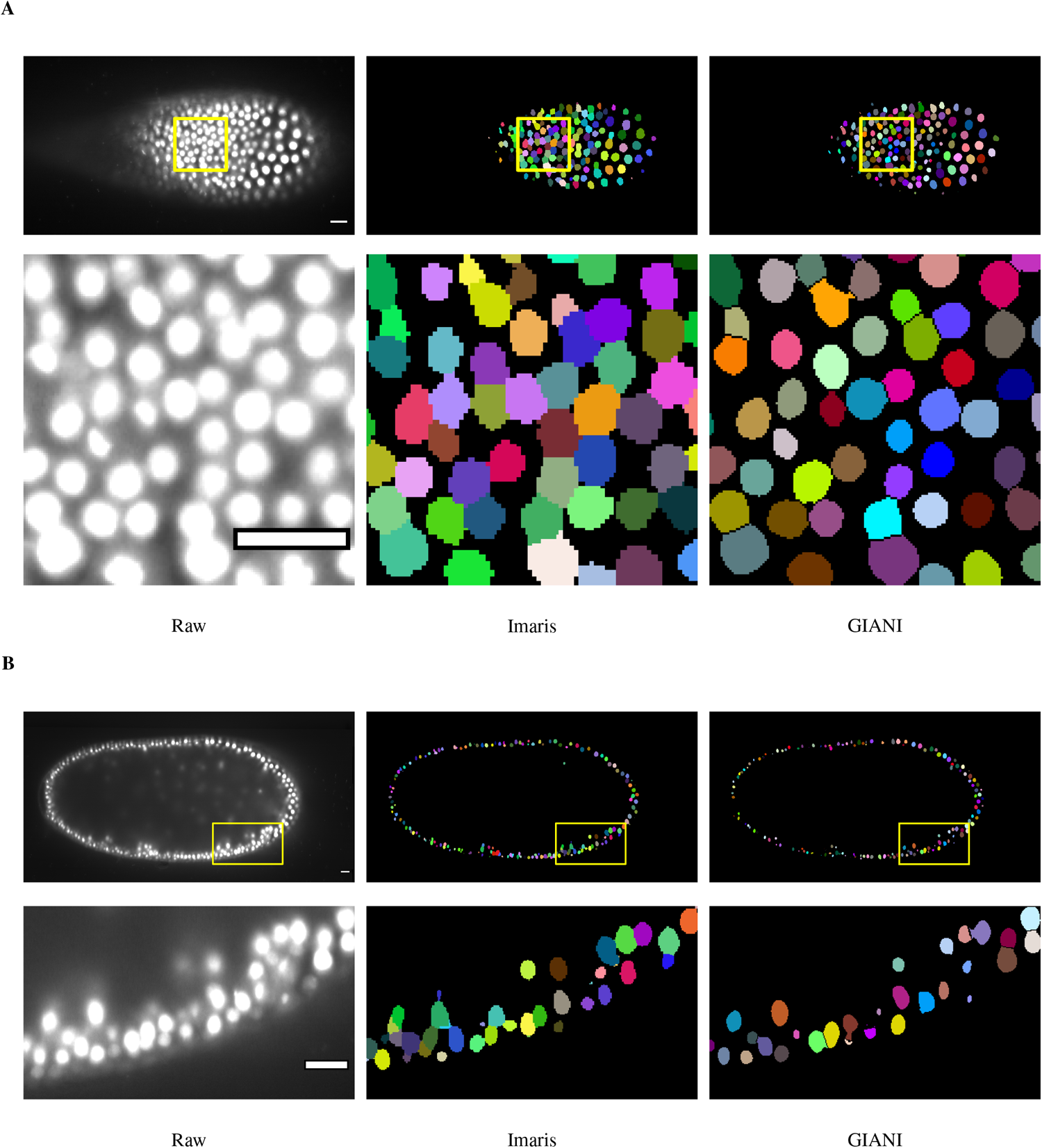
Demonstration of GIANI on a large light sheet microscopy dataset. Illustrations of the segmentations produced by GIANI and Imaris on a *Tribolium castaneum* embryo dataset derived from light sheet microscopy (File S2). Two different slices of the 3D volumes are shown, at approximately 46 *µ*m (A) and 247 *µ*m (B), to illustrate the variation in nuclei morphologies at different depths. In each case, the top row shows the relevant slice of each 3D volume, while the bottom row shows the magnified views of the boxes in the top row images. Approximately 5,600 nuclei were detected in the full volume. Scale bars are all equivalent to 20 *µ*m. The GIANI and Imaris settings used to produce this data are provided as File S6.

### Future developments

Given the ongoing interest in studying cells in “native” 3D extracellular environments [35], future extension to the capabilities of GIANI will include the ability to analyse time-lapse data, as there is currently a lack of open source tools for the quantification of 3D cell migration [36]. The incorporation of additional functionality from TrackMate (and MaMuT [37] or Mastodon) will be explored to facilitate 3D cell tracking.

Future development will also include the replacement of the 3D ROI Manager interface for visualizing results with a new, custom-built interface. At present, segmented objects are saved solely as FIJI ROI files, but it is intended that support for other formats (such as .stl, .ply, .obj and .x3d) will be added to allow the import of objects into a variety of different software.

More generally, with a view to improving and optimising performance, further use of ImgLib2 [38] will be incorporated in future releases. Taking advantage of GPU acceleration is also an aim, most likely by exploiting CLIJ [39]. At present, the analysis of a single simulated embryo used in this study (approx. 2.1 GB) takes between approximately 30 and 60 minutes, depending on the number of CPUs available (Fig S4).

## Conclusion

We have used GIANI to quantitatively analyse mouse embryos in 3D. This analysis has revealed differences in morphology and protein expression between different experimental conditions. Analysis of simulated ground truth data was used to confirm the validity of these results. Further development of GIANI is planned, with the specific aim of improving segmentations in noisy and dense fields of view, common in 3D images of cells. Extension to timelapse analysis is also planned. GIANI is freely available on GitHub (github.com/djpbarry/Giani) and we anticipate that it will be a useful resource for the community to perform routine, automated quantification of complex imaging data.

## Supporting information

**Fig S1.**
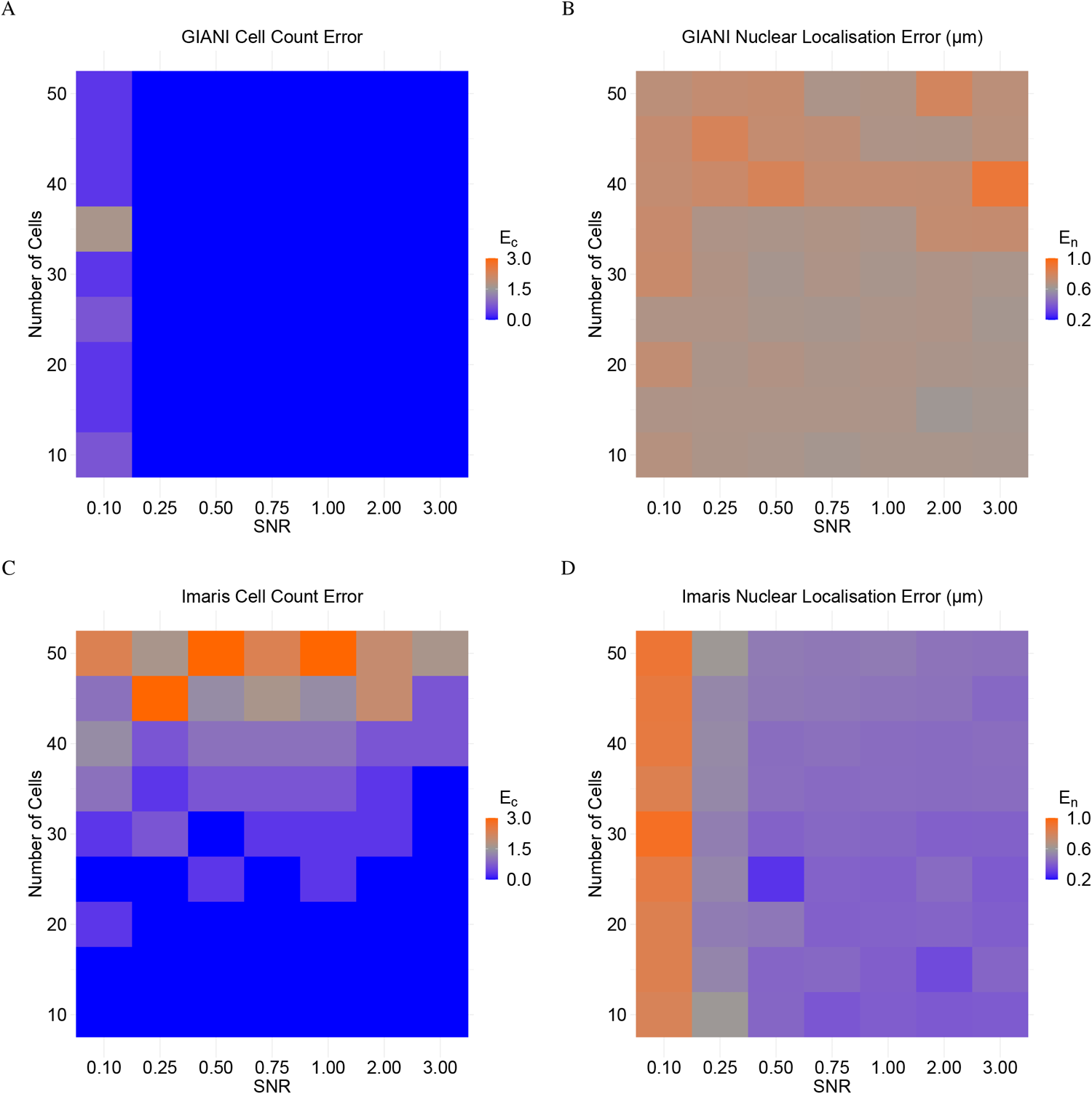
Accuracy of simulated nuclei detection and localisation by GIANI and Imaris. In each of the heat maps, a single tile represents the average of three simulated embryos. The GIANI and Imaris settings used to produce this data are provided as File S7. A: Absolute errors in cell counts (*E*_*c*_, calculated according to Eq 1) produced by GIANI for simulated embryos with the indicated number of cells and signal-to-noise ratios (SNR). B: Absolute errors in nuclear centroid localisation (*E*_*nl*_) produced by GIANI. C: Absolute errors in cell counts produced by Imaris. D: Absolute errors in nuclear centroid localisation produced by Imaris.

**Fig S2.**
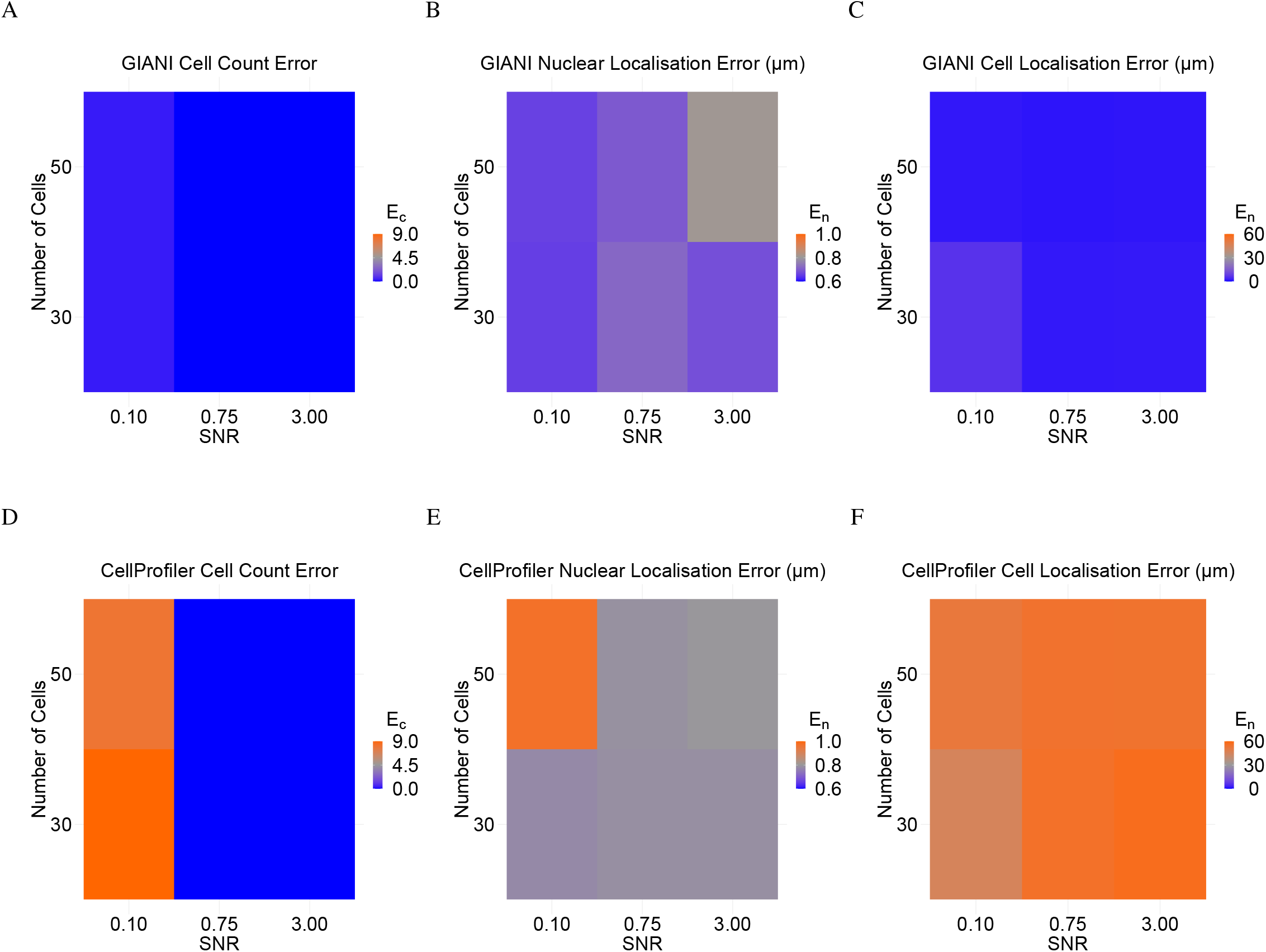
Accuracy of simulated nuclei and cell detection and localisation by GIANI and CellProfiler compared. The data shown in A - C is a subset of that shown in Fig 2F - H. In D - F, each tile represents a single simulated embryo, which was reduced in size to 512 × 512 × 112 voxels prior to running the pipeline. A: Absolute errors in cell counts (*E*_*c*_, calculated according to Eq 1) produced by GIANI for simulated embryos with the indicated number of cells and signal-to-noise ratios (SNR). B: Absolute errors in nuclear centroid localisation (*E*_*nl*_) produced by GIANI. C: Absolute errors in cell centroid localisation error (*E*_*cl*_) produced by GIANI. D: Absolute errors in cell counts produced by CellProfiler. E: Absolute errors in nuclear centroid localisation produced by CellProfiler. F: Absolute errors in cell centroid localisation error produced by CellProfiler. The CellProfiler pipeline used and raw data are provided in File S8.

**Fig S3.**
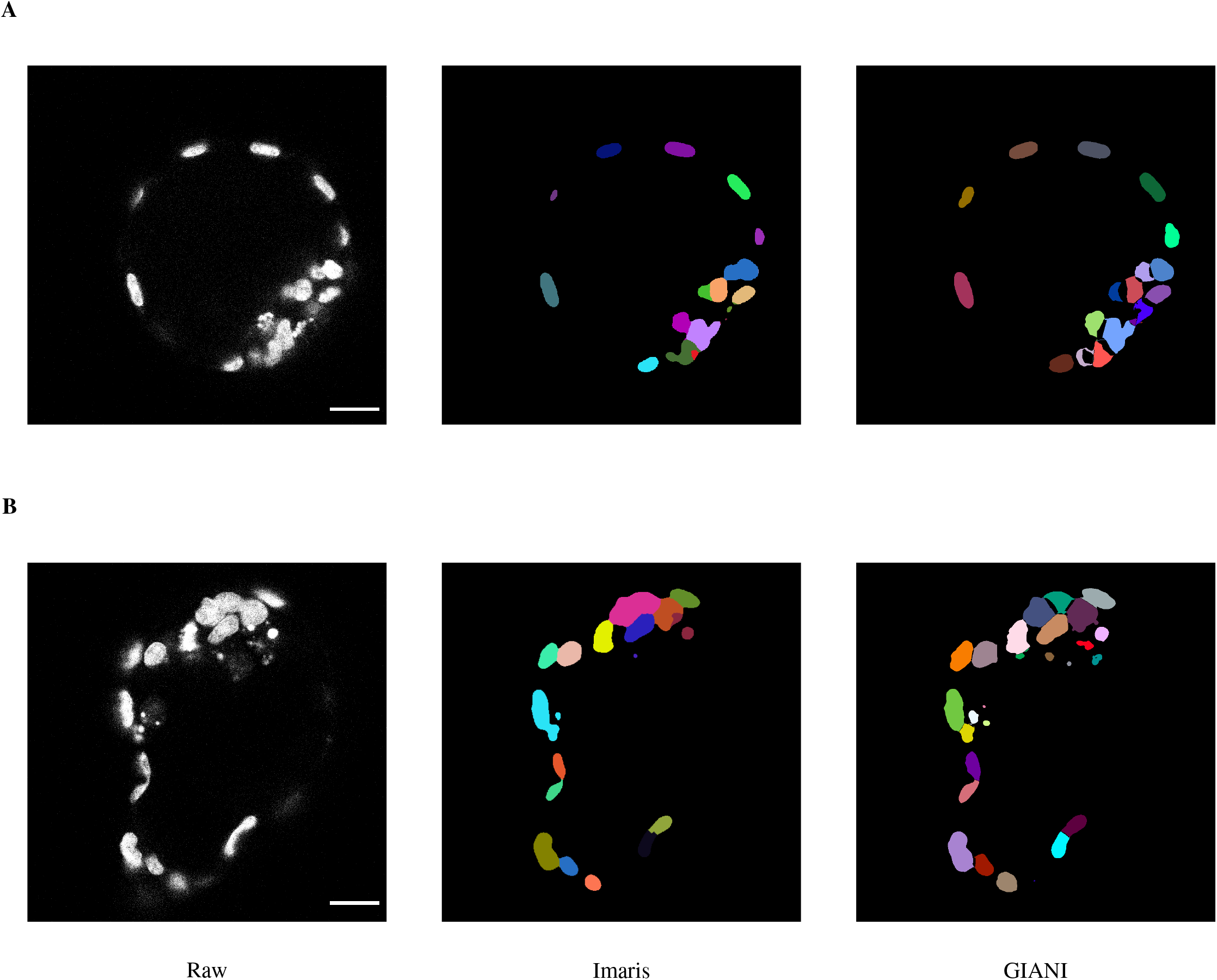
Comparison of mouse blastocyst nuclear segmentations between GIANI and Imaris. A single slice of (A) expanded and (B) hatching blastocysts are shown, together with the segmentations produced by Imaris and GIANI. Scale bars are all equivalent to 20 *µ*m. The GIANI and Imaris settings used to produce this data, together with the raw image data, are provided as File S9.

**Fig S4.**
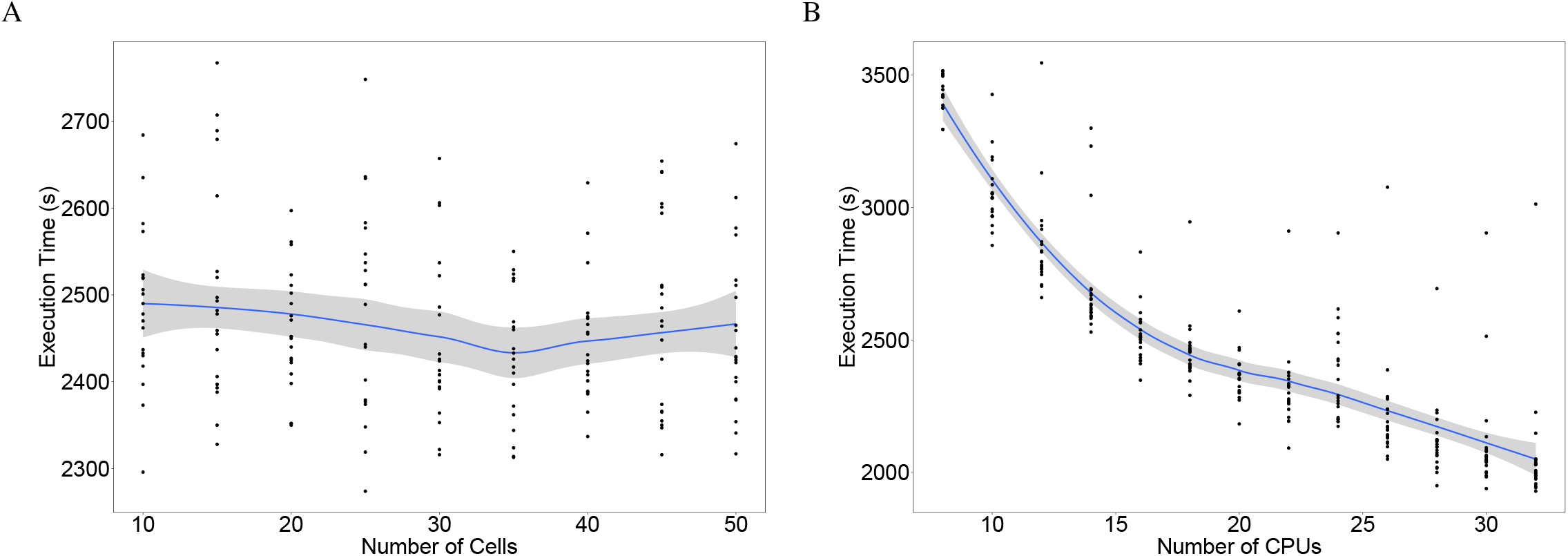
Execution time of GIANI is independent of cell number, but decreases with increasing CPU availability. A: The length of time taken by GIANI to analyse simulated embryos versus the number of cells in the embryo. Each point represents the execution time for a single embryo - approximately 20 embryos were analysed for each cell number. The blue line represents a moving average calculated with LOESS smoothing. The grey bands represent the 95% confidence interval. B: The length of time taken by GIANI to analyse simulated embryos consisting of 30 cells versus the number of available CPUs. Each point represents the execution time for a single embryo - approximately 20 embryos were analysed for each CPU number. The blue line represents a moving average calculated with LOESS smoothing. The grey bands represent the 95% confidence interval.

**File S1 Large simulated image of nuclei**. There are approximately 2000 nuclei in total, each being approximately 10 *µ*m long and 7.5 *µ*m wide. The stack dimensions are 1500 × 1500 × 300 voxels, with each voxel being 0.4 × 0.4 × 2.0 *µ*m, and a total data size of 2.5 GB. Available to download from: https://dx.doi.org/10.5281/zenodo.5270244

**File S2 *Tribolium castaneum* embryo dataset derived from light sheet microscopy**. The dataset is freely available on the Cell Tracking Challenge website (http://celltrackingchallenge.net/3d-datasets) - only the first timepoint was used in the present study. The stack dimensions are 965 × 1871 × 991 voxels, with each voxel being 0.381 × 0.381 × 0.381 *µ*m, and a total data size of 3.3 GB. Available to download from: https://dx.doi.org/10.5281/zenodo.5270323

**File S3 GIANI settings used to produce the data in Fig 2**

**File S4 GIANI settings used to produce the data in Fig 3**

**File S5 GIANI and Imaris settings used to produce the data in Fig 4**

**File S6 GIANI and Imaris settings used to produce the data in Fig 5**

**File S7 GIANI and Imaris settings used to produce the data in Fig S1**

**File S8 CellProfiler pipeline and data used to produce Fig S2**. Available to download from: https://dx.doi.org/10.5281/zenodo.5286507

**File S9 GIANI and Imaris settings and data used to produce Fig S3**. Available to download from: https://dx.doi.org/10.5281/zenodo.5286670

## Acknowledgments

Thanks to Kurt Anderson for his constructive comments on the manuscript. This work was supported by the Francis Crick Institute, which receives its core funding from Cancer Research UK (FC001120, FC001999), the UK Medical Research Council (FC001120, FC001999) and the Wellcome Trust (FC001120, FC001999).

